# Length regulation of multiple flagella that self-assemble from a shared pool of components

**DOI:** 10.1101/436360

**Authors:** Thomas G. Fai, Lishibanya Mohapatra, Jane Kondev, Ariel Amir

## Abstract

Control of organelle size is a problem that has intrigued cell biologists for at least a century. The single-celled green algae *Chlamydomonas reinhardtii* with its two 2agella has proved to be a very useful model organism for studies of size control. Numerous experiments have identi1ed motor-driven transport of tubulin to the growing ends of microtubules at the tip of the 2agella as the key component of the machinery responsible for controlling their length. Here we consider a model of 2agellar length control whose key assumption is that proteins responsible for the intra2agellar transport (IFT) of tubulin are present in limiting amounts. We show that this limiting-pool assumption and simple reasoning based on the law of mass action leads to an inverse relationship between the rate at which a 2agellum grows and its length, which has been observed experimentally, and has been shown theoretically to provide a mechanism for length control. Experiments in which one of the two 2agella are severed have revealed the coupled nature of the growth dynamics of the two 2agella, and we extend our length-control model to two 2agella by considering different mechanisms of their coupling. We describe which coupling mechanisms are capable of reproducing the observed dynamics in severing experiments, and why some that have been proposed previously are not. Within our theoretical framework we conclude that if tubulin and IFT proteins are freely exchanged between 2agella simultaneous length control is not possible if the disassembly rate is constant. However, if disassembly depends on the concentration of IFT proteins at the tip of the 2agellum, simultaneous length control can be achieved. Finally, we make quantitative predictions for experiments that could test this model.

## Introduction

The size regulation of cellular organelles is a fundamental problem in biology ***Marshall (2016)***; ***Milo and Phillips (2015)***. For example, nuclear size is tightly coupled with cell size across a wide range of species ***Hara and Merten (2015)*** and the loss of this coupling mechanism is implicated in various types of cancer ***Zink et al. (2004)***.

A striking example of organelle size control in eukaryotes is the single-celled algae *Chlamydomonas reinhardtii* (***Figure 1(a)***), which uses two flagella to move through its aqueous environment. The lengths of these flagella are tightly controlled; mutants with longer flagella have decreased swimming velocities and beat frequencies ***Khona et al. (2013)*** compared to wild type cells and mutants with unequal flagellar lengths are observed to spin around in circles ***Tam et al. (2003)***.

**Figure 1.**
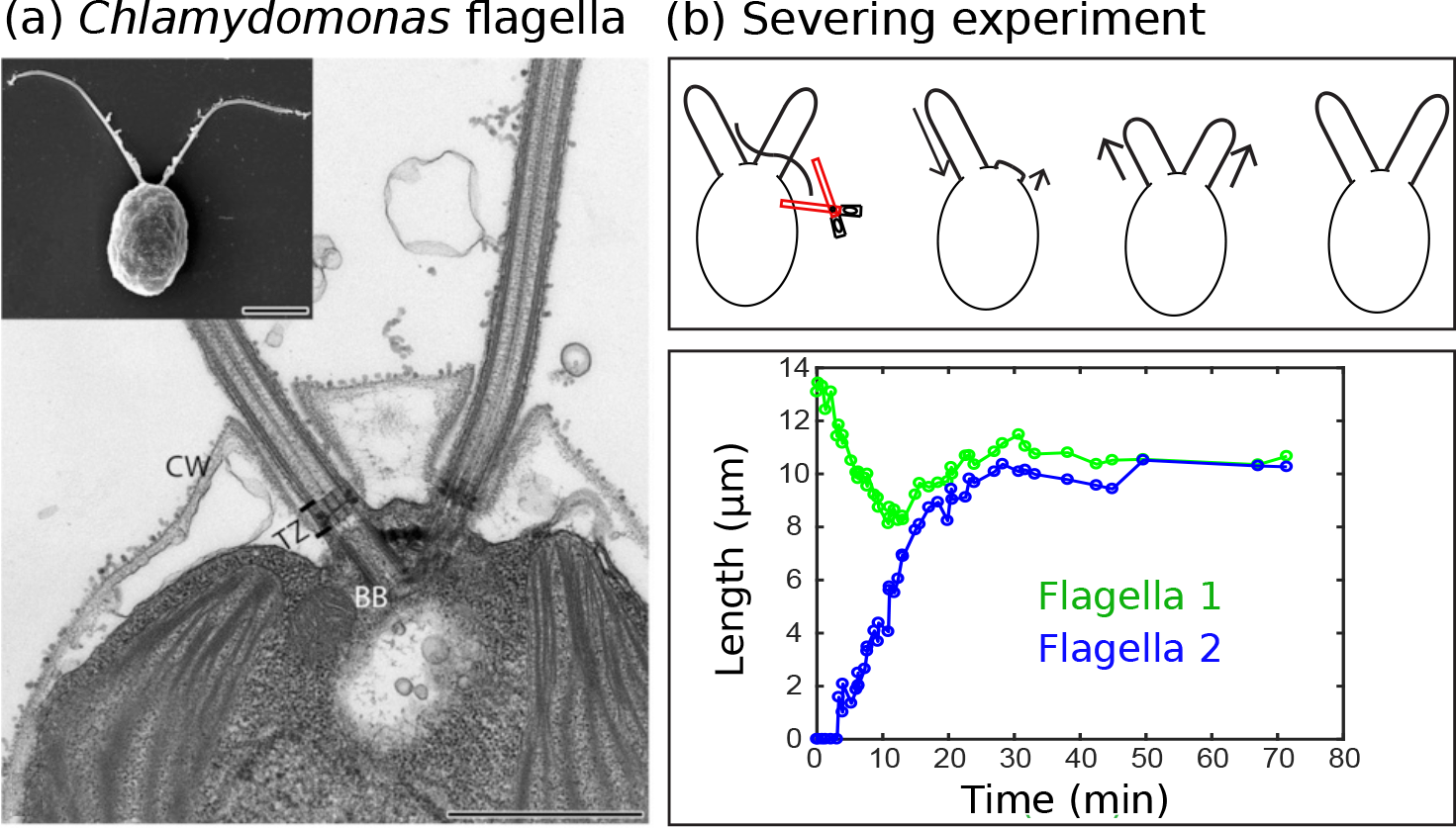
(a) Electron microscopy images of the biflagellate green algae *Chlamydomonas* and its flagella captured by Elisa Vannuccini and Pietro Lupetti (University of Siena, Italy) and reproduced from ***Morga and Bastin (2013)*** (Creative Commons Attribution License CC BY). The inset shows the whole organism (scale bar 5 μm) and the close-up shows the flagellar basal body (BB), transition zone (TZ), and cell wall (CW) (scale bar 1 μm). (b) Severing experiments: after one flagellum is severed, the two flagella equalize at a shorter length and then grow together to the original steady-state length. This is demonstrated in experimental data from ***Ludington et al. (2012)*** courtesy of William Ludington.

It is known from experiments on *Chlamydomonas **Rosenbaum et al. (1969)*** that (i) its two flagella reach steady-state lengths of about 10 μm, and that (ii) the flagellar lengths are correlated since, if one flagellum is severed, the remaining flagellum shortens until it reaches the length of the growing, previously severed flagellum (***Figure 1(b)***). This latter protocol is typically referred to as a “severing” or “long-zero” experiment.

A key process contributing to the assembly of flagella is the continual transport of proteins from the flagellar base to tip and back. The original evidence for this intraflagellar transport (IFT) was provided around 25 years ago by experimental observations in *Chlamydomonas* of particles moving processively along the flagellum at constant speed ***Kozminksi et al. (1993)***. In the time since, a significant body of work has emerged with the goal of understanding the many proteins and biochemical pathways that coordinate this complex process in *Chlamydomonas* and other organisms such as *C. elegans*, as described in several review articles ***Prevo et al. (2017)***; ***Scholey (2008***, 2003); ***Cole (2003)***; ***Rosenbaum et al. (1999)***; ***Rosenbaum and Witman (2002)***; ***Rosenbaum et al. (1999)***.

Already at the time of its discovery, IFT was hypothesized to play a role in flagellar length control by transporting proteins required for assembly at the flagellar tip ***Kozminksi et al. (1993)***. Subsequent work has showed that the proteins in the flagellum are continually exchanged with those in the basal body (***Figure 1(a)***) ***Song and Dentler (2001)***; ***Marshall and Rosenbaum (2001)***; ***Buisson et al. (2013)***. IFT particles containing tubulin are transported along the flagellum by two different motor proteins: kinesin-2 transports IFT particles from the flagellar base to tip (the anterograde direction) whereas dynein carries IFT particles from the tip to the base (the retrograde direction).

These observations motivated the development of mathematical models of flagellar length dynamics such as the balance point model ***Marshall and Rosenbaum (2001)***; ***Marshall et al. (2005)***. In the balance point model, the steady-state flagellar length is achieved when the assembly and disassembly processes, which are in continual competition, come into a balance set by the finite pool of tubulin shared between the flagellum and basal body. The balance point model yields a length-dependent assembly rate by assuming a constant number of transport complexes moving along the flagellum. This is consistent with experimental evidence that the flagellar assembly rate decreases with length ***Rosenbaum and Child (1967)***; ***Marshall and Rosenbaum (2001)***. Notably, however, the balance-point does not explain how the cell could maintain a constant number of transport complexes on the flagellum. Identifying the underlying mechanism of length control remains an open problem, and a subsequent theoretical studies proposed several potential mechanisms that give rise to length-dependent assembly rates ***Ludington et al. (2015)***; ***Hendel et al. (2018)***.

In this work, we propose a simple theoretical framework for the dynamics of flagellar assembly based on the assumption that one or more IFT proteins are present in a limiting amount. Our model captures the essential features of IFT, namely the transport of cargo by molecular motors, and it includes the balance point model as a limiting case. For concreteness, we will assume in our presentation that the rate-limiting protein is a molecular motor; however, the resulting formulas are valid for any rate-limiting IFT protein, as made precise in the Discussion.

We perform a detailed theoretical analysis of the resulting equations in the case of two flagella, considering various possible scenarios for how the finite pools of molecular motors and tubulin may be shared. Whereas a single flagellum reaches a well-defined steady state given finite pools of motors and tubulin, we find that if all pools of biomolecules are shared between multiple flagella and the disassembly rate is length-independent, length control is not achieved regardless of the specific transport mechanisms or model parameters.

To recover simultaneous length control of multiple flagella in the context of a constant disassembly rate, we first explore models in which either the motors or tubulin are shared between flagella, but not both. Supported by experimental observations of microtubule-depolymerizing proteins ***Piao et al. (2009)***; ***Luo et al. (2011)***; ***Hilton et al. (2013)***, we subsequently consider a model in which the disassembly rate is dependent on the local concentration of a depolymerizing IFT protein. Overall, our exploration results in five viable models of flagellar length control (***Figure 6)*** that each generalize to the case of more than 2 flagella while maintaining their salient features.

Four of the five viable models require that the flagella have separate pools for at least one biomolecule. Such a barrier to exchange seems incompatible with the biological evidence of diffusive coupling between basal bodies. It is possible, however, that some proteins are exchanged while others (e.g. tubulin or motors) are not. The fifth viable model, which includes length-dependent depolymerization as an essential ingredient (caused by diffusion and transport within the flagellum of a depolymerizing factor such as kinesin-13 ***Piao et al. (2009)***), allows for all biomolecules to be exchanged between flagella. Depending on the diffusion constant, the assembly rate in this fifth viable model may scale with inverse length as reported in the experimental literature ***Marshall and Rosenbaum (2001)***. In the Discussion, we comment further on the merits of the viable models given the currently available experimental data.

For the five viable models analyzed in this study, we consider potential experimental methods of model validation and/or discrimination. Many cellular processes, including intraflagellar transport, are inherently noisy, and the fluctuations in flagellar length contain valuable information about the underlying biomolecular processes ***Amir and Balaban (2018)***. The essential model parameters may be inferred from measurements of the steady-state length and the fluctuations about steady-state. We use Monte Carlo simulations to estimate the resolution and quantity of time series data that is required for statistically robust data fitting. These estimates may provide useful guidelines for future experiments.

## Results

### Single flagellum

The key biochemical variables represented in our model are the tubulin dimers that form the flagellar microtubules and the molecular motors kinesin and dynein that transport IFT particles from the base to the tip and back. IFT particles combine in the basal body with kinesin, tubulin and dynein to form a complex that is injected into the flagellum ***Cole et al. (1998)***. See the schematic of ***Figure 2***(a).

**Figure 2.**
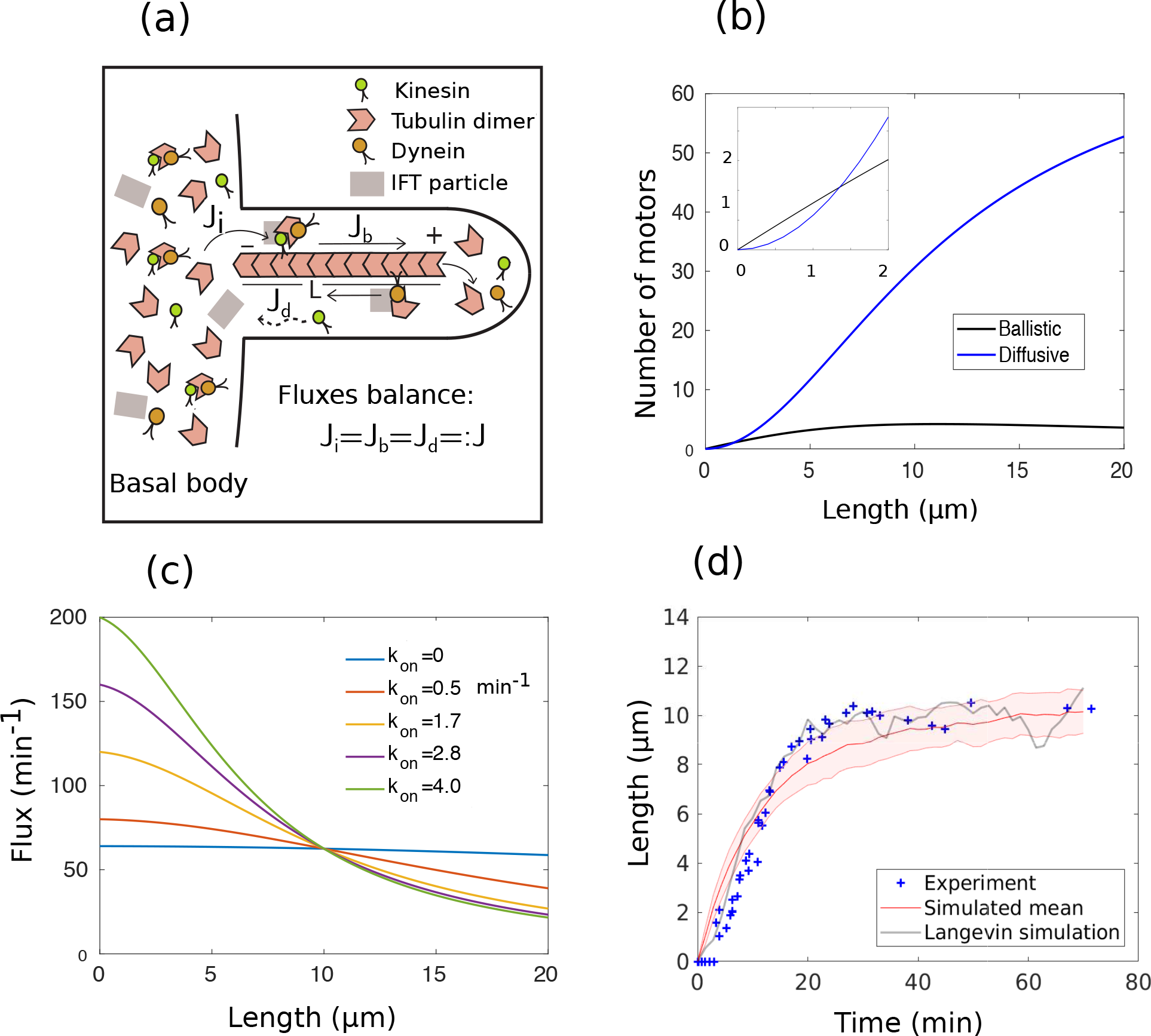
Single flagellar growth dynamics in *Chlamydomonas* in the case of diffusive return of kinesin. (a) IFT particles assemble with kinesin, dynein, and tubulin in the basal body and are injected into the flagellum. In steady-state, the fluxes balance and the injection flux *J*_*i*_, ballistic IFT flux *J*_*b*_, and diffusive IFT flux *J*_*d*_ are all equal. (b) Numbers of molecular motors undergoing ballistic and diffusive motion on the flagellum at a given length, obtained using the parameter values in ***Table 1***. There is a crossover at *L* ≈ 1.5 μm (inset) after which diffusive motion is dominant. (c) IFT particle flux vs. length for several choices of the on-rate of molecular motors *k*_on_ (see (3)). (d) Model simulations show that an initially severed flagellum grows until it reaches steady-state, consistent with the behavior observed experimentally in ***Ludington et al. (2012)***. The shaded red area illustrates the mean and standard deviation computed from 50 realizations of the Langevin equation (see (29) in Materials and methods).

We assume that the injection of IFT particles into the flagellum is rate-limited by the supply of molecular motors. The total number of molecular motors is assumed to be conserved, and we consider the case that the rate-limiting motor undergoes ballistic motion only (as for dynein ***Toropova et al. (2017)***) and the case that the rate-limiting motor undergoes ballistic transport in the anterograde direction and diffusive motion in the retrograde direction (as for kinesin ***Chien et al. (2017)***). The total amount of tubulin is also conserved in our model. The flagellar assembly rate, i.e. the rate of addition of tubulin dimers to the microtubule ends at the tip of the flagellum, is assumed proportional to the flux of IFT particles from the basal body into the flagellum times the amount of free tubulin in the basal body. This assumption is a consequence of mass-action kinetics of IFT particle assembly. (Note that we consider the cell volume to be fixed, in which case the amount and concentration of a given biomolecule may be used interchangeably.)

The ballistic and diffusive IFT dynamics are fast compared to changes in length. The timescale of flagellar length dynamics e.g. the recovery time after severing is of the order of 10 minutes, whereas the molecular motors involved in IFT take at most tens of seconds to traverse the length of the flagellum by either ballistic or diffusive motion. Based on this separation of timescales, we follow ***Hendel et al. (2018)*** by neglecting the time delay between injection and tubulin delivery and by treating IFT as a quasi-steady state process.

We use a stochastic differential equation to model the flagellar length dynamics *L*(*t*). We are motivated to include stochasticity because of experimental observations of flagellar length fluctuations and since measuring fluctuations about steady-state can be used to discriminate between candidate models. Our stochastic model follows ***Wemmer and Marshall (2018)***, a combined experimental and theoretical study that illustrates how noise in the flagellar length control system affects biological function. Whereas the theoretical component of this previous study focused on the balance point model, here we include fluctuations within a more general theoretical framework. We model the growth of a *single* flagellum by

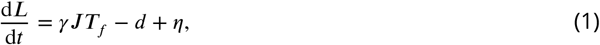

where *J* is the flux of IFT particles (to be determined), *T*_*f*_ is the amount of free tubulin in the basal body, *d* is the disassembly rate (assumed constant for now), and *γ* is a constant. To model the molecular noise due to the inherent stochasticity of binding and unbinding events, we add a noise term *η*. Mathematically, it represents white noise forcing of magnitude ϒ, i.e. *η* is a random variable with autocorrelation 〈*η*(*t*)*η*(*t*′)〉 = ϒ^2^*δ*(*t* − *t*′). See the explanation preceding (29) in Materials and methods for details.

In the context of our model, the quasi steady-state assumption implies an injection rate *J* that exactly balances the ballistic and diffusive fluxes. As shown in (33) and (42) of Materials and methods, the fluxes satisfy

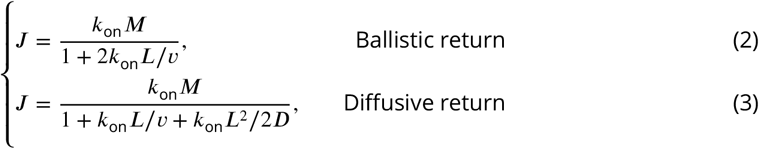

where *k*_on_ is the rate constant of motor injection, *M* is the total number of motors, *v* is the speed of ballistically-moving motors in IFT, and *D* is the diffusion constant of motors in the flagellum (with parameter values given in ***Table 1***). Note that in (3) and throughout the manuscript, terms written in the form *x*/*yz* are shorthand for *x*/(*yz*), e.g. *k*_on_*L*^2^/2*D* is to be interpreted as *k*_on_*L*^2^/(2*D*).

**Table 1.** Parameter values and de1nitions

| Symbol | Definition | Value | Units | References |
| --- | --- | --- | --- | --- |
| **Parameters** |  |  |  |  |
| $L_{ss}$ | Steady-state length | 10–12 | $\mu\text{m}$ | *Marshall and Rosenbaum (2001); Rosenbaum et al. (1969)* |
| $T/N$ | Tubulin pool per flagellum | 38–47 | $\mu\text{m}$ | *Marshall et al. (2005)* |
| $d$ | Disassembly rate | 0.5 | $\mu\text{m}/\text{min}$ | *Marshall and Rosenbaum (2001); Ludington et al. (2012)* |
| $v$ | IFT speed | 2.5–3 | $\mu\text{m}/\text{s}$ | *Kozminksi et al. (1993); Buisson et al. (2013)* |
| $D$ | Diffusion coefficient | 1.7 | $\mu\text{m}^2/\text{s}$ | *Chien et al. (2017)* |
| $\gamma k_{\text{on}} M/N$ | Assembly rate per tubulin | $2.3 \times 10^{-2}$–$3.6 \times 10^{-2}$ | $\text{min}^{-1}$ | Fit |
| $k_{\text{on}}$ | Injection rate constant | 0.8–4.5 | $\text{min}^{-1}$ | Fit |
| $\gamma$ | Prefactor in (1) | $2.5 \times 10^{-4}$ | | Estimate (**Appendix 1**) |
| **Variables** |  |  |  |  |
| $N$ | Number of flagella | | | |
| $T_f$ | Free tubulin | | $\mu\text{m}$ | |
| $M$ | Total motors | | | |
| $M_f$ | Free motors | | | |
| $M_b$ | Ballistic motors | | | |
| $M_d$ | Diffusing motors | | | |
| $J$ | Flux | | $\text{min}^{-1}$ | |
| $c_d(x)$ | Motor concentration | | $\mu\text{m}^{-1}$ | |
| $\bar{c}_d$ | Average concentration | | $\mu\text{m}^{-1}$ | |

Near the steady-state length in the case of diffusive return, we find that most motors on the flagellum undergo diffusive motion, although at very small lengths ballistic motion is dominant (***Figure 2***(b)). The flux decreases with length (***Figure 2***(c)). On a qualitative level, this decreasing trend is supported by experimental measurements ***Dentler (2005)***, although more data is needed to make a quantitative comparison. In the appropriate limits, this expression recovers existing theories in which the length dependence goes purely as 1/*L* or 1/*L*^2^. In the case of ballistic return, or for diffusive return in the limit *D ≫ Lv*, the 1/*L* scaling of ***Marshall and Rosenbaum (2001)*** is recovered. For diffusive return in the limit *D ≪ Lv*, the 1/*L*^2^ scaling of ***Hendel et al. (2018)*** is recovered. A distinction between our formula for the flux and those in the aforementioned paper is the presence of the constant term in the denominator, which follows from injection according to the law of mass action. Consequently, the flux does not become singular at *L* = 0 in our formulation. It is interesting to note that the flux has a similar functional form to the familiar substrate production rate in Michaelis-Menten enzyme kinetics ***Fall (2002)***; this is because of the separation of timescales assumption invoked in both derivations.

Substituting the above expressions for the flux into the growth rate equation (1) results in

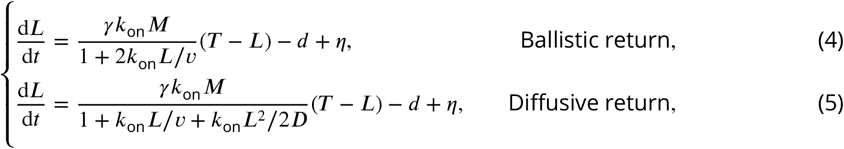

where *T*_*f*_ = *T* − *L* by conservation of the total tubulin pool *T*. Each of the above models is consistent with the results of flagellum severing experiments, i.e. when the severing experiment is simulated by setting the initial length to zero, the length increases until it reaches its steady-state *L*_*ss*_ (***Figure 2***(d)).

In the case of ballistic return, the steady-state length is given by

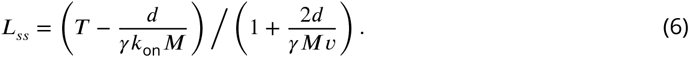

In the case of diffusive return, solving (5) for the steady-state results in a quadratic equation for *L*_*ss*_. One root is always negative, leaving the solution

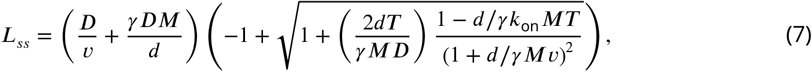

which is positive provided that *T > d*/*γkM*. If *T* ≤ *d*/*γkM*, the flagellar length shrinks to zero. (We reiterate the convention that terms such as *d*/*γk*_on_*MT* and *d*/*γMv* are to be interpreted as *d*/(*γk*_on_*MT*) and *d*/(*γMv*), respectively.)

Expanding to first order in Δ*L* := *L* − *L*_*ss*_, we find that the steady-state is stable, i.e. d(Δ*L*)/d*t* = −*λ*(Δ*L*) with *λ* > 0 given in (36) of Materials and methods. Based on the parameters estimated in ***Appendix 1*** for diffusive return, the associated timescale *τ* := 1/*λ* is approximately 15 min, which is consistent with experiment. This timescale is long compared to the few tens of seconds needed for molecular motors to traverse the flagellum in IFT, which justifies *a posteriori* our approximation of IFT as a quasi-steady state process. We next consider the parameter space associated with the length dynamics. Introducing the nondimensional length 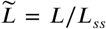, time 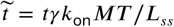, and random forcing 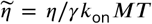 we may rewrite (4) and (5) in terms of the nondimensional parameters *π*_1_ = *d*/*γk*_on_*MT*, *π*_2_ = *L*_*ss*_/*T*, *π*_3_ = *k*_on_*L*_*ss*_/*v*, and 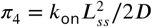 as

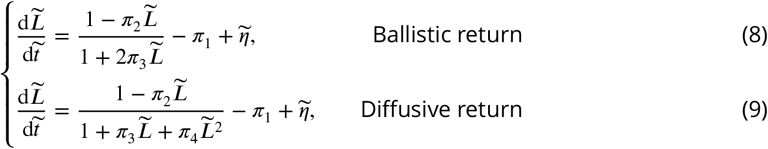

We may interpret these numbers as follows: *π*_1_ is the ratio of disassembly and assembly rates, *π*_2_ is the fraction of the tubulin pool taken up by the flagellum at steady-state, *π*_3_ = *τ*_*b*_/*τ*_*i*_ is the ratio of the ballistic timescale *τ*_*b*_ := *L*_*ss*_/*v* of IFT transport to the timescale 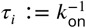 of injection, and *π*_4_ = *τ*_*d*_ /*τ*_*i*_ is an analogous ratio involving the diffusive timescale 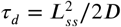. (We could equivalently think of *π*_3_ and *π*_4_ as ratios of lengthscales related to the same physical processes.)

We next show that, in a special limit, the length dynamics with diffusive return are nearly equivalent to the model of ***Hendel et al. (2018)***. In terms of the experimentally measured parameters and those estimated in ***Appendix 1***, we find *π*_1_ ≈ 0.4, *π*_2_ ≈ 0.2, *π*_3_ ≈ 0.1, and *π*_4_ ≈ 0.8. The relatively small values of *π*_2_ and *π*_3_ suggest that it may be useful to consider the limit *π*_2_ → 0 (i.e. no tubulin depletion) and *π*_3_ → 0 (i.e. instantaneous ballistic motion). In this limit, we have

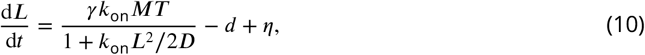

recovering the model of ***Hendel et al. (2018)*** with the only distinction that, as mentioned above, in our model there is an additional constant term in the denominator. In this limit, the steady-state length satisfies

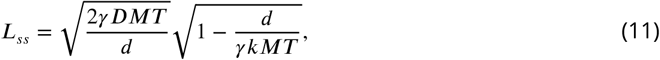

which carries an extra factor of 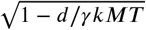 in comparison with the formula of ***Hendel et al. (2018)***.

### Two flagella

In the case of two flagella with lengths *L*_1_(*t*) and *L*_2_(*t*) the length dynamics are given by

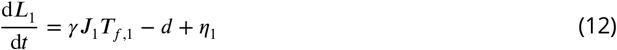

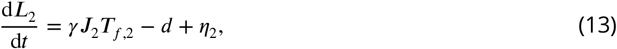

where *J*_*i*_ and *T*_*f,i*_ for *i* = 1, 2 denote the fluxes and free amounts of tubulin for the two flagella, and *η*_1_ and *η*_2_ are independent white noise forcing. Severing experiments illustrate that the flagella are coupled. We consider various modes of coupling, illustrated in ***Figure 3***(a), that give rise within our model to different forms of *J*_*i*_ and *T*_*f,i*_ and investigate their consequences for length control by focusing on the stability of solutions to the steady-state deterministic equations

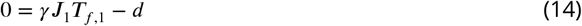

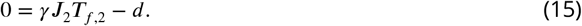

**Figure 3.**
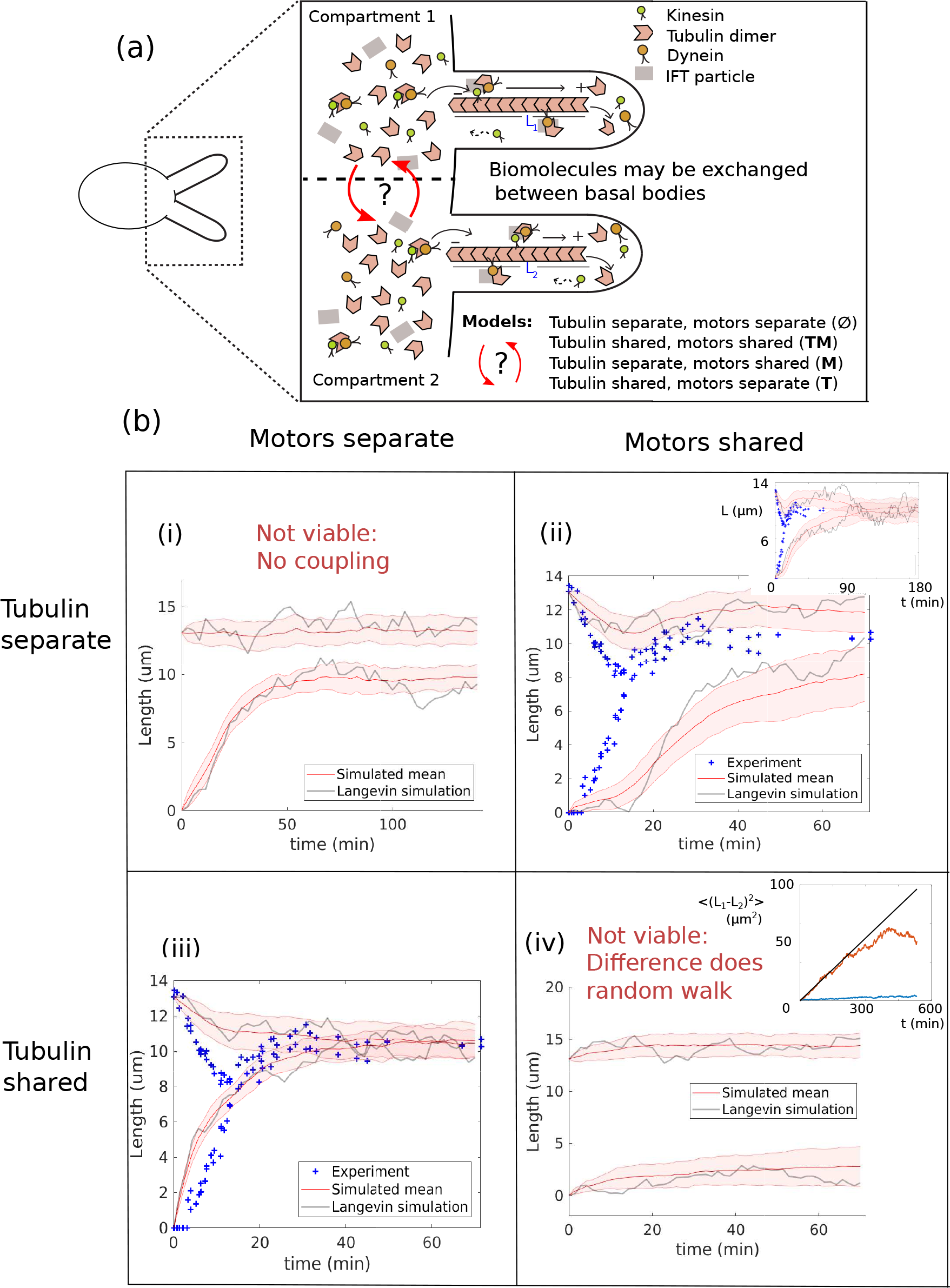
Length dynamics of two flagella assembling from shared pool of building blocks. The case of diffusive motor return is shown; similar results are obtained with ballistic return. (a) Flagellar assembly in *Chlamydomonas reinhardtii* and modes of coupling between basal bodies. (b) Simulations of severing experiment by models **ø**, **M**, **T**, and **TM**. (i) In the case of separate pools of both motors and tubulin, the unsevered flagellum does not decrease in length. (ii) When only motors are shared, the flagella reach equal steady-state lengths (inset shows longer times). (iii) When only tubulin is shared, the two flagellar lengths reach a shared steady-state on a timescale consistent with experiment. (iv) When both tubulin and motors are shared, the lengths do not equalize. The difference between the two flagellar lengths undergoes a random walk, as shown by the mean square displacement (inset, average over 50 runs. The deviation from linearity is caused by the non-negativity of flagellar lengths, which limits the maximum difference.) In all the above simulations, to capture the loss of the tubulin from severing and the gradual recovery by intracellular regulation mechanisms, we initialize the tubulin pool with 14 μm less of tubulin and linearly increase it over the first 20 minutes to reach the steady-state value of *T*.

#### Tubulin separate, motors separate (ø)

The case of individual tubulin pools and individual motor pools leads to two uncoupled instances of (5), and therefore following the logic of the single flagellum case yields steady state lengths given by (7). While these uncoupled equations trivially yield simultaneous length control, they are fundamentally inconsistent with the coupling observed between flagella in severing experiments, and in particular the significant decrease in the length of the unsevered flagellum (***Figure 3***(b)(i)). Therefore we exclude this model as a viable candidate.

#### Tubulin separate, motors shared (M)

When motors are shared through a common pool, IFT particles are injected into either flagellum with equal probability. Therefore *J*_1_ = *J*_2_ =: *J* and it can be shown by a straightforward generalization of (2) and (3) (see Model (**M**) in Materials and methods) that the flux satis1es

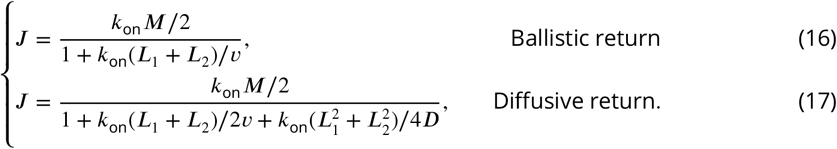

The model yields simultaneous length control (***Figure 3***(b)(ii)). Because the tubulin pools are separate the steady-state equations are given by

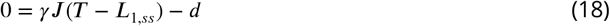

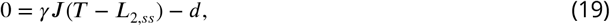

and it follows that *L*_1*,ss*_ = *L*_2,*ss*_ after subtracting the above equations. With *M* rescaled to *M*/2, the steady-state equations are identical to the corresponding steady-state equations (4) and (5) for a single flagellum. Therefore the steady-state lengths for ballistic and diffusive return satisfy (6) and (7), respectively, upon rescaling *M* → *M*/2. See Model (**M**) in Materials and methods for detailed derivations. The stability of the steady-state solution is proven in ***Appendix 2*** in the context of diffusive return. (The proof for ballistic return is essentially identical.)

Given the large amount of turnover and interaction between different cellular compartments, the assumption of separate tubulin pools in this model is questionable. There is experimental evidence that tubulin is exchanged between each flagellum and its basal body ***Song and Dentler (2001)***; ***Marshall and Rosenbaum (2001)***; ***Buisson et al. (2013)***. However, we cannot rule out the possibility of separate tubulin pools, as we are not aware of experimental evidence of tubulin exchange between flagella. Therefore we retain this as a viable model.

#### Tubulin shared, motors separate (T)

We next consider the case in which tubulin is shared but the motor pools are separate. The separate motor pools yield decoupled fluxes so that the flux is nearly identical to that of model **ø**. However, in this case the equations are coupled through the shared tubulin pool term *T* − *L*_1_ − *L*_2_. The model equations (see (75)–(76) in Materials and methods) have a similar form to existing models ***Marshall et al.(2005)***; ***Hendel et al. (2018)***, in which the assembly rates involve a factor of *T* − *L*_1_ − *L*_2_ and either a 1/*L*_*i*_ or 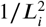-dependence in the denominator, for *i* = 1, 2. In fact, the present model recovers these two dependences on *L*_*i*_ in the appropriate limits, as discussed in Results in the context of a single growing flagellum.

As illustrated in ***Figure 3***(b)(iii), this model yields simultaneous length control and the flagella reach their steady-state lengths on a timescale consistent with experiment. Based on this analysis of the **M** and **T** models, sharing tubulin *or* motors is in principle suZcient to attain simultaneous length control. This is independent of ballistic versus diffusive return of motors to the basal body. As mentioned above, in light of the constant turnover and biochemical interactions between different cellular compartments, the assumption of separate motor pools may be incorrect. Indeed, there is preliminary evidence of IFT protein exchange between flagella based on experiments using photoconvertible fluorophores ***Marshall (2018)***. It is generally believed that both tubulin and motor pools are shared between flagella, and we next consider this case.

#### Tubulin shared, motors shared (TM)

The flux resulting from the shared motor assumption is the same as in (17), and we are left with the steady-state equations

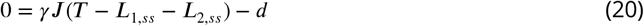

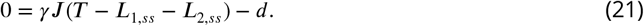

Because the two steady-state equations are identical, there is only a single equation for the two unknowns *L*_1*,ss*_ and *L*_2*,ss*_, which makes the lengths indeterminate. The model in which all biomolecules are shared between flagella does not yield simultaneous length control! There are infinitely many solutions to the equations, none of which is asymptotically stable (since performing a linear stability analysis corresponding yields a zero eigenvalue, as shown in ***Appendix 2)***. The same issue also appears in the dynamics; since d*L*_1_/d*t* = d*L*_2_/d*t* this model cannot possibly explain the simultaneous positive and negative growth rates for the two flagella observed in the severing experiment.

Note that this conclusion is *independent* of the parameters. That is, regardless of whether the transport is diffusive, ballistic, or some combination, the model **TM** with shared tubulin and shared motors with constant disassembly does not yield simultaneous length control of two flagella (***Figure 3***(b)(iv)), and therefore is not a viable model.

The above analysis shows that to obtain length control in our framework under the assumption of constant disassembly, either tubulin or motors may be shared, but not both. However, the two viable models **M** and **T** identified above are incompatible with the widely-held belief that biomolecules are continually exchanged between flagella. It follows that either the conventional wisdom of shared pools is wrong or something is missing from the space of models considered. This motivates us to extend our study beyond the shared pool models considered so far.

#### Tubulin shared, motors shared and concentration-dependent disassembly (TM*)

We next consider a model that allows for full biomolecule exchange and replaces the constant disassembly assumption with a concentration-dependent disassembly rate. The assumption of a constant rate of disassembly was based on experiments on mutants in which IFT was disabled ***Marshall et al. (2005)***. However, these experiments do not preclude motor-dependent disassembly since they were performed in the absence of motor dynamics. Indeed, subsequent experiments in organisms with intact IFT led to 50-fold greater disassembly rates than those measured in the absence of IFT ***Ludington et al.(2012)***.

Experimental observations that some kinesin species (e.g. kinesin-13) participate in microtubule disassembly ***Piao et al. (2009)*** provide a potential biochemical basis for IFT-dependent disassembly, and in what follows we take the disassembly rate to depend on motor concentration. We replace the constant disassembly by a disassembly rate of the form *d*_0_ + *d*_1_*c*_*d*_ (*L*), where *c*_*d*_ (*L*) is the concentration of diffusing motors at the tip of the flagellum. In general the disassembly rate may be a complicated function of concentration, and in that case the model is only appropriate near steady-state, where the first-order term in the Taylor expansion dominates.

Under the assumption of ballistic return this model does not yield length control; diffusive return is required to generate a non-constant concentration along the length of the flagellum. Assuming diffusive return the flux and concentration at the flagellar tip are related by *c*_*d*_ (*L*) = *J L*/*D* (see (37) in Materials and methods), we may rewrite the governing equations as

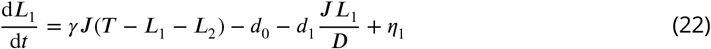

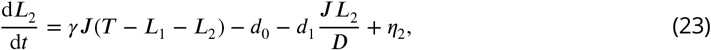

where as before in the case of shared motors

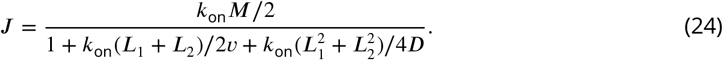

The concentration-dependent disassembly model with shared tubulin and shared motors (**TM\***) exhibits simultaneous length control (***Figure 4)***. Subtracting (23) from (22), it follows immediately that *L*_1*,ss*_ = *L*_2*,ss*_ =: *L*_*ss*_, and solving for the steady-state results in

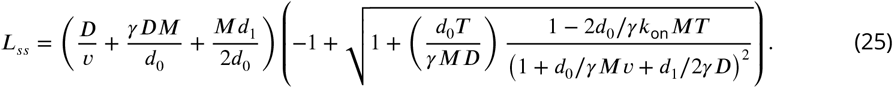

**Figure 4.**
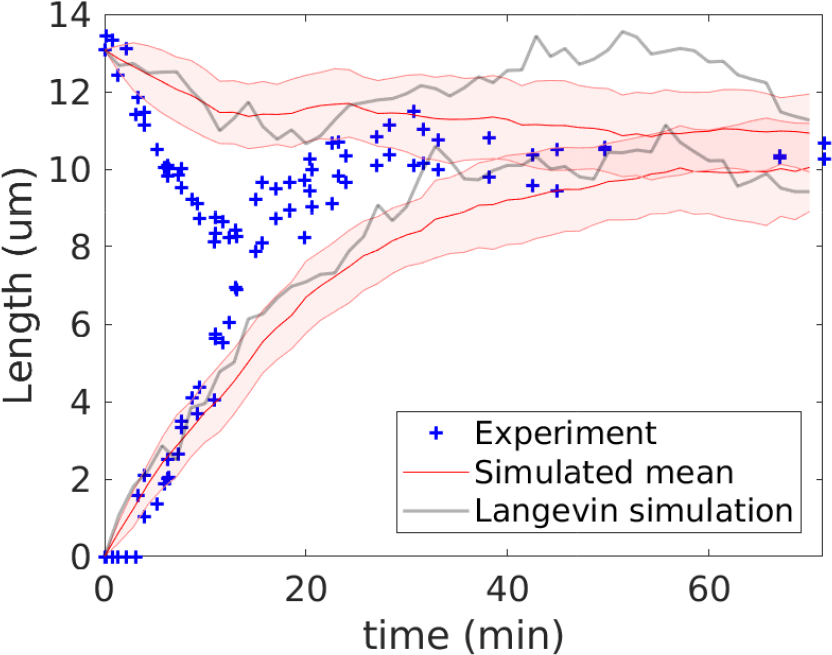
Simultaneous length control is achieved in a model with shared tubulin, shared motors, and a disassembly rate that depends on the concentration of molecular motors. Here we set *k*_on_ = 4.5 min^−1^, *d*_0_ = 0.1 μm/min, and *d*_1_ is computed by (25) to obtain the observed steady-state length.

In the Appendix it is shown that this solution is stable. Therefore, concentration-dependent disassembly allows for simultaneous length control when all biomolecules are shared between flagella, and results in a third viable model. Diffusive return is a critical ingredient in model **TM\***; with ballistic return it would not yield simultaneous length control. This is because, as remarked above, the non-constant diffusive motor concentration profile is necessary to make the growth rates of *L*_1_ and *L*_2_ distinct, which provides two independent equations for the steady-state lengths.

### Generalization to *N* > 2 flagella

Taking inspiration from the eukaryotic parasite Giardia, which has four pairs of flagella ***Dawson and House (2010)***; ***McInally and Dawson (2016)***, we demonstrate the simultaneous length control attained for *N* = 8 flagella (***Figure 5)***.

We may generalize the concentration-dependent disassembly model **TM\*** to arbitrary flagellar number *N*. (The other viable models **T** and **M** may be generalized in a similar manner.) Because motors are shared, the injection fluxes are equal so that *J*_*i*_ = *J* for all *i* = 1 … *N*, with *J* satisfying

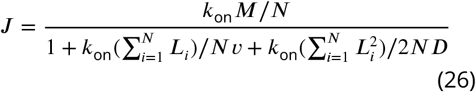

and length dynamics given by

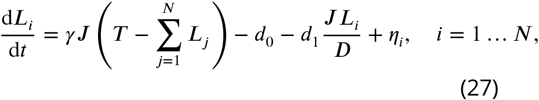

**Figure 5.**
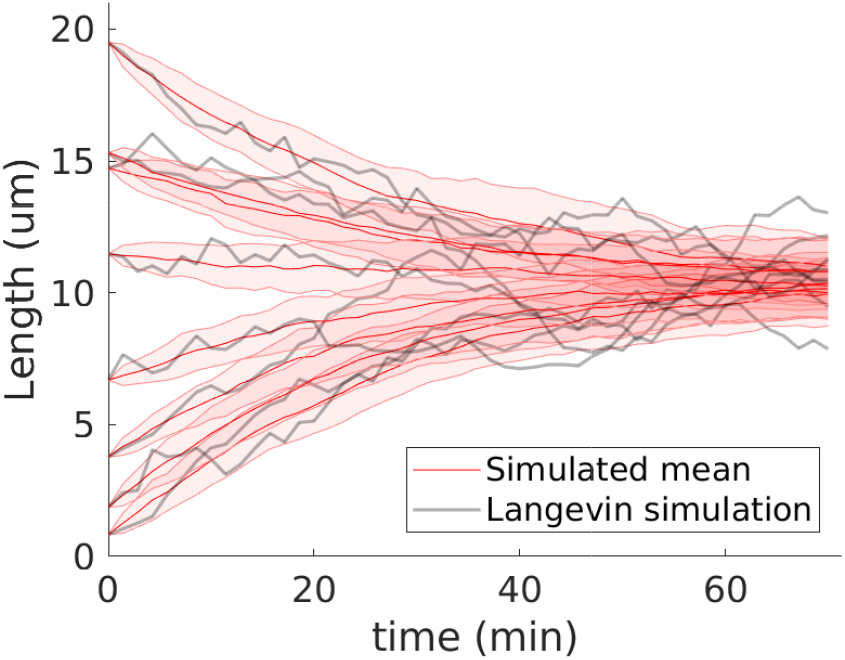
The viable models generalize to arbitrary flagellar number *N*. We demonstrate using the concentration-dependent disassembly model **TM\*** with *N* = 8 flagella and the larger shared pool *T* = 336, *d*_0_ = 0.1, and *d*_1_ = 120; otherwise all parameters are as in the biflagellate case.

Taking any pairwise difference between the *i*^th^ and *j*^th^ equations at steady-state yields immediately *L*_*i,ss*_ = *L*_*j,ss*_, so that the steady-state lengths are equal to

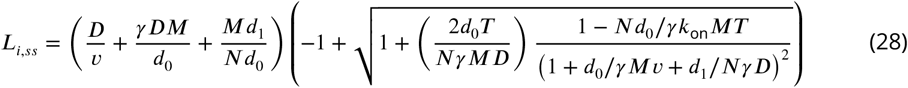

for all *i* = 1…*N*. Stability follows from analyzing the linearized equations, as shown in ***Appendix 2***.

## Discussion

In this work, we explore models of flagellar length dynamics in the context of organelle size regulation. We capture the essential aspects of IFT relevant to length control, i.e. the motor-driven transport of tubulin across the flagellum. As in ***Hendel et al. (2018)***, since the timescale of transport by IFT is a few seconds, whereas changes in flagellar length take place on a timescale of a few minutes, we treat IFT as a quasi-steady-state process in which the anterograde and retrograde fluxes of motors are balanced.

It has been well-established that the biomolecules represented by our model, i.e. tubulin and molecular motors, are recycled within each flagellum and its basal body. There is also the possibility of turnover *between* flagella. We have used our theoretical model to investigate the consequences of sharing pools of tubulin and/or motors between flagella on their length dynamics. We take the assembly rate to be proportional to the flux of IFT particles times the amount of free tubulin. In our initial exploration, we take the disassembly rate to be constant. We find that sharing both tubulin and motors leads to an indeterminate system of equations regardless of the details of the model, whereas sharing either tubulin or motors, but not both, results in simultaneous length control of both flagella. Interestingly, the model in which only tubulin is shared (model (**T**)) recovers essential features of the balance point model ***Marshall and Rosenbaum (2001)***, in particular the decay with 1/*L* in the growth rate.

This theoretical exploration illustrates the dramatic consequences in behavior that can occur when biomolecules are shared between compartments, and highlights the importance of knowing which proteins are exchanged between the basal bodies in the context of flagellar length control. In particular, having a protein that is *not* exchanged can provide simultaneous length control by a limited pool mechanism. Future experiments to identify which proteins are exchanged between flagella, e.g. by tagging with photoconvertable fluorophores as in ***Marshall (2018)***, are needed to assess the validity of this mechanism.

We investigated additional models that are consistent with the widely-held belief that biomolecules are exchanged continually between flagella, e.g. in which both the motor and tubulin pools are shared. In particular, we considered the possibility that disassembly is not constant but depends on the local concentration of molecular motors. Because the diffusive gradient is linear at steady-state and the basal body is taken to act as a diffusive sink, the concentration at the flagellar tip depends on length. This leads to simultaneous length control regardless of whether the disassembly rate is strongly or weakly dependent on concentration. We conclude that the title of ***Hendel et al. (2018)*** is apt; the linear concentration gradient generated by diffusion indeed acts as a ruler! However, it cannot function on its own. Diffusion must be combined with a mechanism such as concentration-dependent disassembly; regardless of whether the motor dynamics are ballistic, diffusive, or some combination, sharing all biomolecules does not yield length control within our theoretical frame-work if the disassembly rate is constant. To put it another way, within the space of the five viable models (***Figure 6)***, diffusion is not essential for length control in the **M** and **T** models, whereas it *is* essential for model **TM**\*.

**Figure 6.**
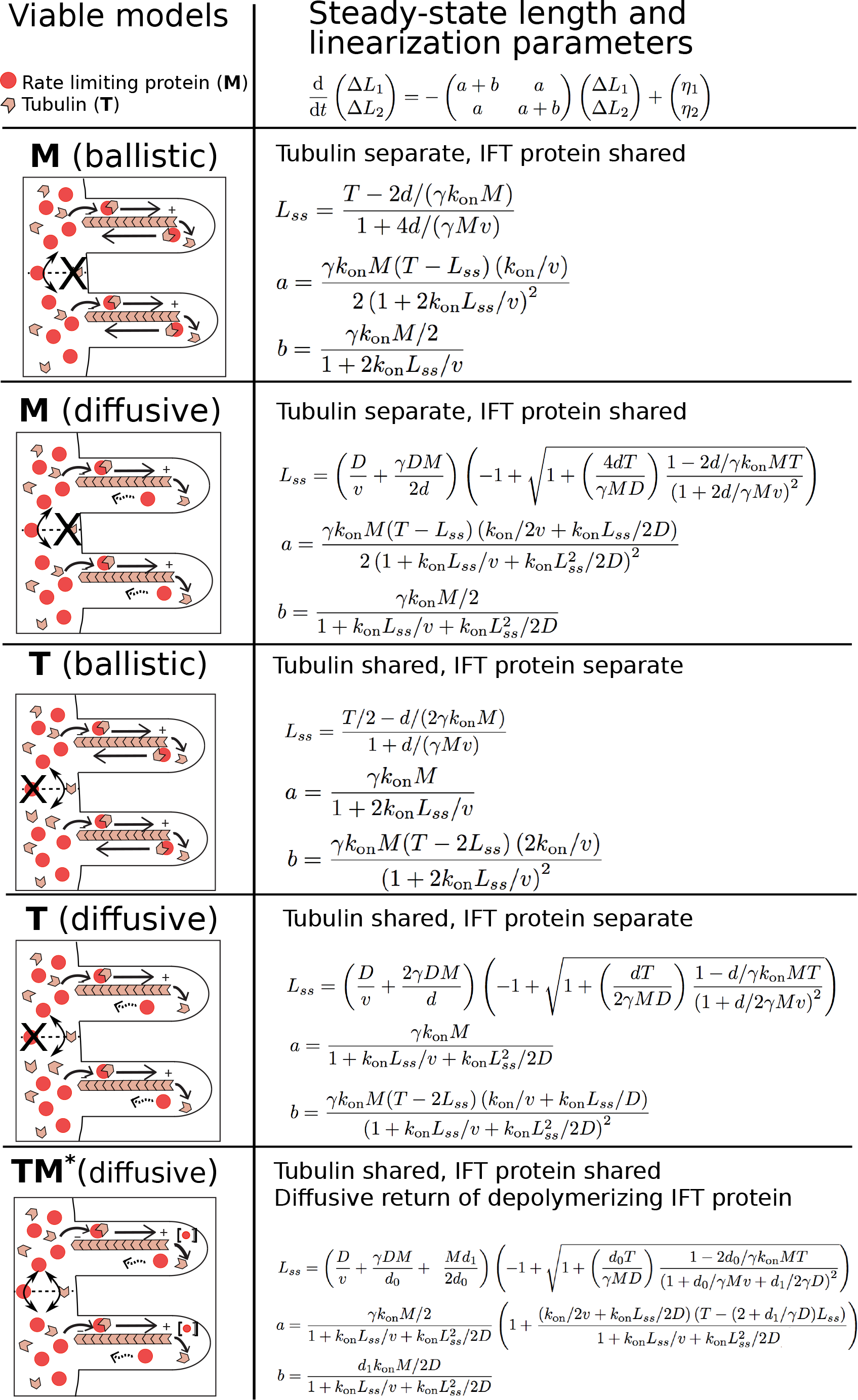
Our theoretical exploration results in five viable models

Note that, although the present discussion has assumed that a molecular motor is the rate-limiting IFT protein, our formulation is actually more general. If a non-motor IFT protein is rate-limiting for injection, the same considerations and resulting formulas apply with the number of motors replaced by the rate-limiting protein number and accounting for the relevant mode of transport (i.e. ballistic or diffusive return). This abstracted view of the rate-limiting protein is represented schematically in ***Figure 6***.

Diffusion is an essential ingredient for length control in model **TM**\*, and depending on the diffusion constant the growth rate decays as either 1/*L* or 1/*L*^2^. Using the diffusion constant measured for kinesin results in a 1/*L*^2^ decay; although the experimentally-determined growth rate was originally 1t to a curve with 1/*L* decay ***Marshall et al. (2005)***, we find that fitting the same data to a curve with 1/*L*^2^ decay or to the second-order polynomial decay of model **TM**\* lead to 1ts of comparable quality.

Among the viable models discussed here, only model **TM**\* is consistent with both full biomolecule exchange and simultaneous length control. This model requires a rate-limiting IFT protein coupled to microtubule disassemby, shared between flagella, and diffusing back from the flagellar tip. Further experiments are needed to establish whether a protein meeting these criteria exists. One promising candidate is aurora-like kinase CALK, which has been shown to influence disassembly through its state of phosphorylation ***Luo et al. (2011)***; ***Cao et al. (2013)***. Another promising candidate is CNK2, a NIMA-related protein kinase known to localize to flagella ***Bradley and Quarmby (2005)***. It has been shown that a long-flagella Chlamydomonas mutant lacking a functional CNK2 exhibits decreased disassembly rates ***Hilton et al. (2013)***, so that this protein would also be compatible with the concentration-dependent disassembly assumption. Future experiments will determine whether flagellar lengths are controlled by the diffusion of CALK, CNK2, or some other protein, or whether additional physical mechanisms beyond those included in this theoretical framework are required to explain the observed phenomena.

On the side of theory, a promising avenue to discriminate between the potential models is by studying length fluctuations about steady-state. The fluctuation spectra of each model provides a signature that can be used to assess both the general model framework and to test the autocorrelation timescales predicted by each model ***Amir and Balaban (2018)***. In ***Appendix 3*** we have described the spectra predicted by the viable models. We have mimicked experiments by performing stochastic simulations with various sampling rates and durations to provide constraints on the experimental parameters necessary to obtain robust and statistically significant results through an analysis of fluctuations. We find that while significant amounts of data are needed in order to extract information from fluctuations, the sampling intervals and time durations needed to obtain 10% error are within the realm of experimental plausibility (see ***Appendix 3***).

## Materials and methods

We use a stochastic ordinary differential equation model for the flagellar length dynamics. The key biochemical variables represented in our model are the tubulin monomers that form the flagellar axoneme and the molecular motor that carries IFT particles along the flagellum.

### Single flagellum

To introduce the model in a simpler setting, we consider first the case of a single flagellum with time-dependent length *L*(*t*). In our model the flagellar assembly rate is proportional to the flux *J* of IFT particles times the size of an IFT particle, which is in turn proportional to the amount of free tubulin *T*_*f*_ in the basal body. In other words, once the flux of IFT motors has been established, forming a type of conveyor belt, tubulin dimers have a constant probability per unit time of joining with a motor, i.e. of jumping onto the conveyor belt. (Equivalently, one may think of a sequence of biochemical reactions: first, IFT particles are generated at a rate *J*, and second, tubulin forms a complex with IFT at a rate proportional to *JT*_*f*_.) We take the disassembly rate to be equal to a constant *d* for now. Finally, to account for random fluctuations we include a white noise forcing term *η* with autocorrelation 〈*η*(*t*)*η*(*t*′)〉 = |ϒ|^2^*δ*(*t* − *t*′), where *δ*(*t* − *t*′) represents the Dirac delta function. This yields a Langevin equation for the growth rate:

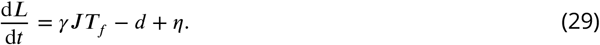

We assume that the rate of injection of IFT particles into the flagellum is rate-limited by the supply of molecular motors. The molecular motors are assumed to be conserved, and we consider two cases separately: first the case in which the rate-limiting IFT motor undergoes ballistic motion in both directions on the flagellum, and second the case in which the motor undergoes ballistic motion from the flagellar base to tip and diffusive motion in the retrograde direction. Note that, although there is evidence that IFT loading occurs in trains of various sizes ***Wren et al. (2013)***; ***Ludington et al.(2013)***, we have not included differential loading into our model as the simpler model in which dimers load independently is sufficient for agreement with the severing experiments.

#### Ballistic return

Motors are conserved having total number *M* = *M*_*f*_ + *M*_*b*_, where *M*_*f*_ is the number of motors freely available in the basal body and *M*_*b*_ is the number of motors moving ballistically on the flagellum in IFT particles. In our model the flux, or injection rate, is proportional to the number of free molecular motors *M*_*f*_ so that *J* = *k*_on_*M*_*f*_ according to mass action kinetics, with first-order rate constant *k*_on_.

We assume tubulin is conserved with total amount *T* = *T*_*f*_ + *T*_*b*_ + *L*, where *T*_*f*_ is the amount of tubulin freely available in the basal body, *T*_*b*_ is the amount moving ballistically while attached to IFT particles, and *L* is the amount incorporated in the flagellum. As the flagellum grows it incorporates more tubulin and the size of the free tubulin pool decreases. We assume that the amount of tubulin undergoing IFT is negligible, i.e. *T*_*b*_ ≪ *T*_*f*_ + *L*, so that *T* ≈ *T*_*f*_ + *L*. This approximation is justified since it can be shown by the quasi steady-state assumption that *T*_*b*_/*T_f_ <* 2*γk*_on_*M L*_*ss*_/*v*, and consequently *T*_*b*_/*T_f_* < 2.6 × 10^−3^ in terms of the parameters contained in ***Table 1***.

By mass action and conservation of motors, the flux of motors may be expressed as

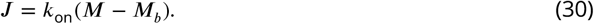

Given velocities *v*_*a*_ and *v*_*r*_ in the anterograde and retrograde directions, respectively, in ballistic motion the flux is related to the corresponding concentrations in a simple manner. The flux satisfies *J* = *c*_*a*_*v*_*a*_ = *c*_*r*_*v*_*r*_, where *c*_*a*_ and *c*_*r*_ are the concentrations of motors moving in the anterograde and retrograde directions. (It follows from the quasi steady-state assumption that the anterograde and retrograde fluxes are balanced, so that there is a single flux *J*.) Therefore

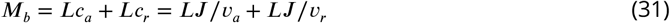

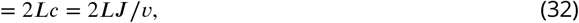

where *c* = (*c*_*a*_ + *c*_*r*_)/2 is the arithmetic mean and *v* = 2(1/*v*_*a*_ + 1/*v*_*r*_)^−1^ is the harmonic mean (i.e. the time-averaged velocity of motion along the flagellum ***Marshall and Rosenbaum (2001)***). Substituting (32) into (30) and solving for the flux results in

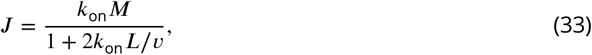

which from (29) and the relation *T*_*f*_ = *T* − *L* yields the growth rate

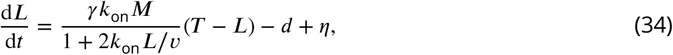

where we neglect time delays and treat IFT as a quasi steady-state process based on the timescale separation discussed earlier between IFT transit and flagellar length dynamics. The steady-state d*L*/d*t* = 0 is reached at

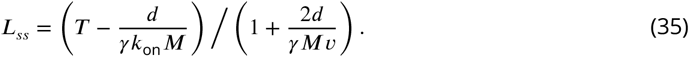

Expanding to first order in Δ*L* = *L* − *L*_*ss*_ yields

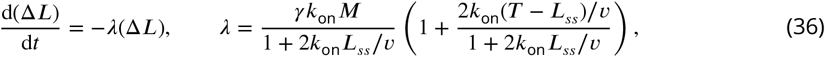

so that the steady-state is stable since *λ* > 0. The steady-state length *L*_*ss*_ is positive provided that *T M > d*/*γk*_on_. This sets a theoretical lower limit on the product of total motors and tubulin needed to observe stable flagella, which makes sense given the constant disassembly term that must be balanced to reach steady-state.

#### Diffusive return

Direct experimental evidence shows that kinesin motors do not move ballistically through all phases of IFT but rather, after ballistic anterograde motion, diffuse back upon reaching the tip of the flagellum ***Chien et al. (2017)***. This mechanism has been considered in a recent theoretical study ***Hendel et al. (2018)***. We next examine how the results above obtained under the assumption of purely ballistic motion change when the rate-limiting IFT protein must return from tip to base by diffusion. In this case, there is an additional variable ***M***_*d*_ that represents the number of motors diffusing from the tip to the base, and ***M*** = ***M***_*f*_ + ***M***_*b*_ + ***M***_*d*_.

We assume that ballistic and diffusive IFT dynamics are fast compared to changes in length and are well-approximated by a quasi-steady state process. This implies the diffusive flux *D∂c*_*d*_ /*∂x* equals the injection rate *J*, and moreover the steady-state condition *D∂*^2^*c*_*d*_ /*∂x*^2^ = 0 implies the concentration profile *c*_*d*_ (*x*) of the diffusing motors is linear, i.e.

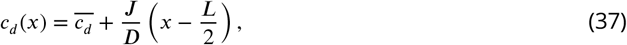

in which 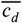 is the average concentration along the flagellum and *D* is the diffusion constant. The flagellar base is taken to act as a diffusive sink so that *c*_*d*_ (0) = 0. This boundary condition implies that ^1^

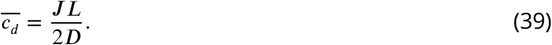

Upon including diffusion, we may replace (30) in the above derivation for the purely-ballistic case by

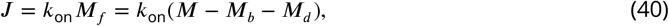

where *M*_*b*_ = *J L*/*v* given diffusive motion from base to tip and moreover

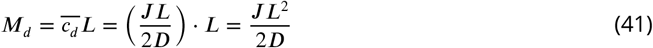

for the diffusively-moving motors. This results in the expression

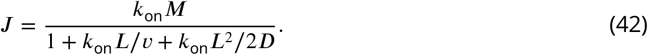

The flux is therefore a quadratic function in length. Substituting this expression for the flux into the growth rate equation (29) results in

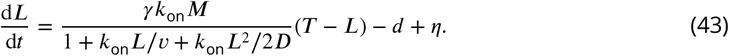

The rate-limited flux arising from a finite number of motors therefore provides a mechanistic derivation that can account for either 1/*L* or 1/*L*^2^-length dependence under different limits of model parameters. Upon introducing the measured parameters and simply computing the terms in (42), we find *k*_on_*L*_*ss*_/*v* ≈ 0.1 and 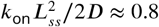, so that diffusion dominates over ballistic motion at steady-state, but none of the terms is negligible over the complete short to long-length dynamics.

Solving (43) for the steady-state results in a quadratic equation for *L*_*ss*_. One root is always negative, leaving the positive solution

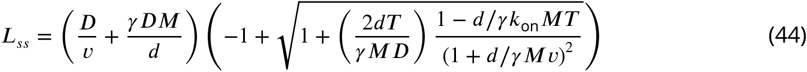

We can evaluate the stability of this solution by linearizing about *L*_*ss*_. Expanding to first order in Δ*L* := *L* − *L*_*ss*_, we find d(Δ*L*)/d*t* = −*λ*(Δ*L*) with *λ* a positive constant given by

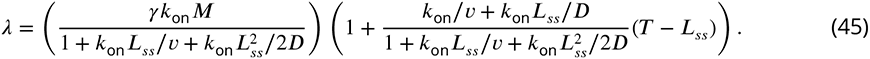

### Two flagella

In the case of two flagella with lengths *L*_1_(*t*) and *L*_2_(*t*) the length dynamics are given by

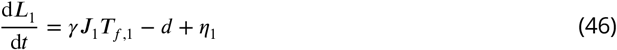

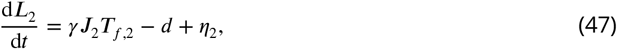

where *J*_*i*_ and *T*_*f,i*_ for *i* = 1, 2 denote the fluxes and free amounts of tubulin for the two flagella, which may be equal if they are shared. The fluctuation terms *η*_1_ and *η*_2_ are taken to be uncorrelated white noise forcing having the covariance

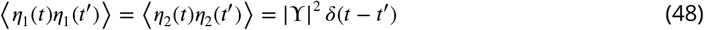

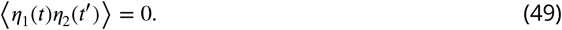

We will consider various modes of coupling between the flagella giving rise to different forms of *J*_*i*_ and *T*_*f,i*_ and their consequences for length control. To assess whether a particular length control model is viable we focus on stability of solutions to the steady-state deterministic equations

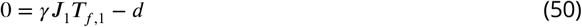

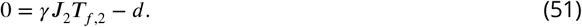

#### Tubulin separate, motors separate (ø)

The presence of individual tubulin pools and individual motor pools leads to two uncoupled instances of the single flagellum dynamics, that is

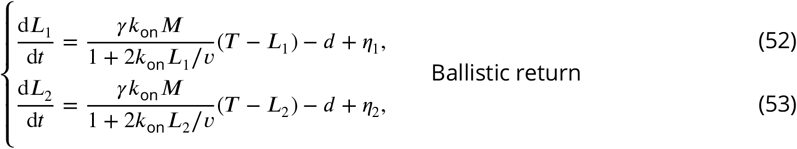

and

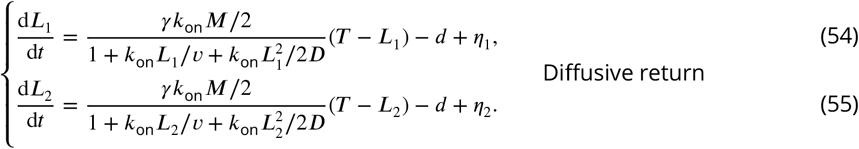

This leads to steady state lengths given by (35) and (44).

#### Tubulin separate, motors shared (**M**)

In the case of individual tubulin pools, we have *T*_*f*,1_ = *T* − *L*_1_ and *T*_*f*, 2_ = *T* − *L*_2_. We next compute the motor flux in the cases of ballistic and diffusive return.

##### Ballistic return

We have

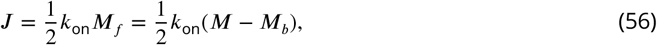

where the factor of one-half comes from assuming equal injection probability into either flagellum. Further, *M*_*b*_ = 2*J* (*L*_1_ + *L*_2_)/*v* so that

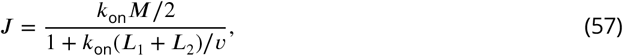

and consequently

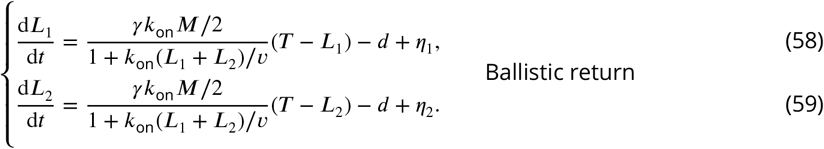

The steady-state length is given by The resulting equations for the steady-state length are

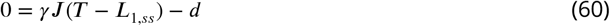

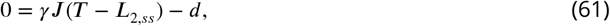

which yield

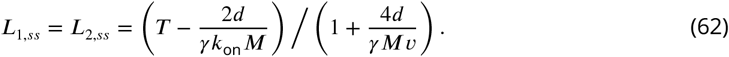

##### Diffusive return

The flux may be calculated according to

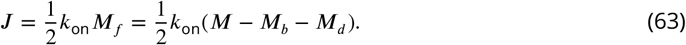

Since the ballistic motion is only in the base to tip direction *M*_*b*_ = *J* (*L*_1_ + *L*_2_)/*v* and

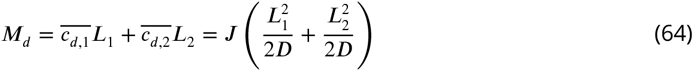

so that

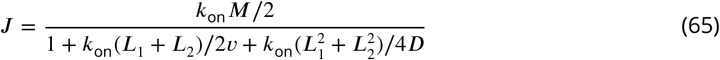

which yields the flagellar length dynamics

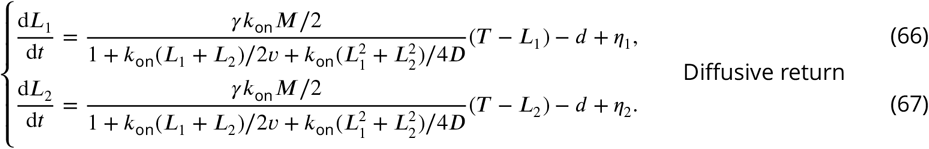

In the case of diffusive return the steady-state solution is

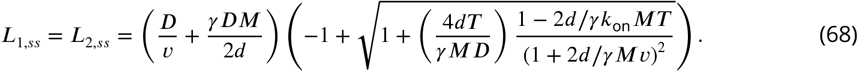

#### Tubulin shared, motors separate (**T**)

We next consider the case in which only tubulin is shared. The separate pools of motors yield decoupled fluxes

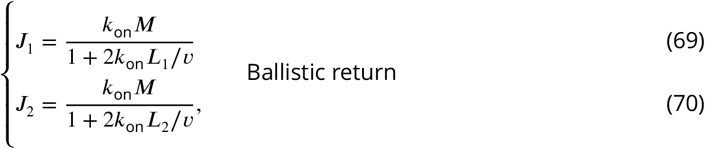

and

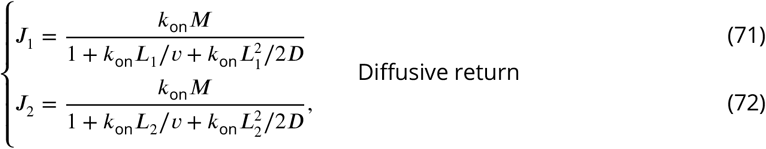

which leads to the systems of equations

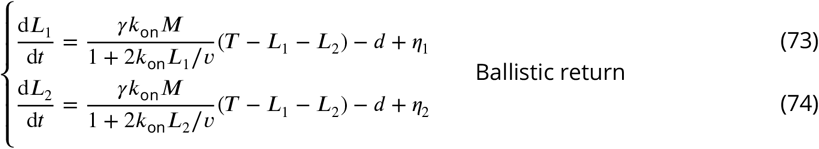

and

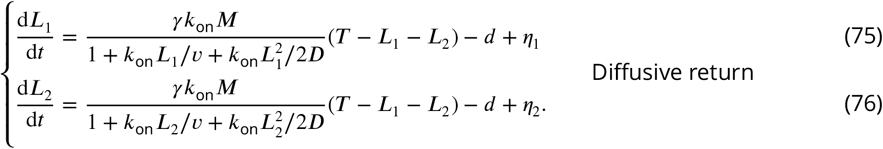

These systems of equations are nearly identical to those of model **ø** given by (52)–(53) and (54)– (55), with the notable exception that the equations are coupled through the shared tubulin pool term *T* − *L*_1_ − *L*_2_. This model yields simultaneous length control. The steady-state lengths satisfy *L*_1*,ss*_ = *L*_2*,ss*_. The resulting steady-state equations are identical to those from (4) and (5) for a single flagellum, with *T* rescaled to *T*/2 and *γ* rescaled to 2*γ*. Therefore the steady-state lengths for ballistic and diffusive return satisfy (6) and (7), respectively, upon rescaling *T* → *T* /2 and *γ* → 2*γ*.

#### Tubulin shared, motors shared (**TM**)

We finally consider the case in which both tubulin and motors are shared through a common pool. By the shared tubulin pool assumption *T*_*f,* 1_ = *T*_*f*, 2_ = *T* − *L*_1_ − *L*_2_ and by the shared motor pool assumption the injection rates satisfy *J*_1_ = *J*_2_ ≡ *J*. Regardless of the particular form of the flux, we are left with the steady-state equations

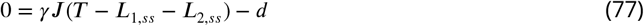

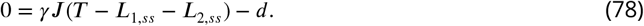

There is only a single equation for the two unknowns *L*_1*,ss*_ and *L*_2*,ss*_ and the lengths are indeterminate. Upon adding random noise, the sum fluctuates around a specific value whereas the difference undergoes a random walk (inset to ***Figure 3***(b)(iv)).

## Acknowledgments

We acknowledge useful discussions with Wallace Marshall, William Ludington, Shashank Shekhar, and Shane Mcinally. We thank Jie Lin and Ethan Levien for reading a draft of this manuscript and offering helpful feedback. We acknowledge funding support from National Science Foundation grant DMS-1502851 (TGF), grant DMR-1610737 (JK), MRSEC-1420382 (JK), the Simons Foundation (JK, LM), the A. P. Sloan Foundation (AA), and the Kavli Institute (AA).

## Appendix 1

### Parameter estimation

To estimate the model parameters, we fit to experimentally-measured data. First, we use the extrapolated data point from ***Marshall et al. (2005)*** that the growth rate upon severing a flagellum is approximately 0.4 μm/min. This implies

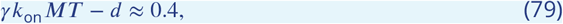

where fluctuations have been neglected. We first estimate the product *γk*_on_*M*. To do so, we use the measured disassembly rate *d* = 0.5 μm/min ***Marshall and Rosenbaum (2001)*** and an estimated instantaneous tubulin pool of *T* = 25-40 μm. This yields *γk*_on_*M* = 2.3 × 10^−2^−3.6 × 10^−2^ min^−1^. The estimate *T* for the tubulin pool available to a single flagellum in the instant after severing is based on the reported amount 76-94 μm for the total tubulin pool shared between two flagella ***Marshall et al. (2005)*** after accounting for some tubulin lost after severing. This estimate for *γk*_on_*M* is not expected to be very precise, particularly given that the total tubulin pool size from ***Marshall et al. (2005)*** was itself obtained by fitting the parameters of a related model to data on mutants with extra flagella.

We can subsequently estimate *k*_on_ by fitting it to the steady-state length observed in the severing experiment. Returning to (43) and once again neglecting fluctuations, we may express *k*_on_ in terms of quantities that have already been measured or estimated:

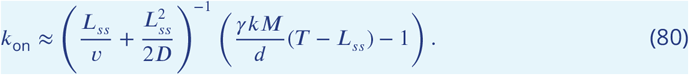

Plugging in the values for *γkM*,*d*, and *T* above as well as the measured values *v* = 150 μm/min ***Kozminksi et al. (1993)***; ***Buisson et al. (2013)***, *D* = 102 μm^2^/min ***Chien et al. (2017)***, and *L*_*ss*_ = 10 μm measured in ***Ludington et al. (2012)***, we obtain *k*_on_ = 0.8–4.5 min^−1^. See the parameter values and definitions in ***Table 1***.

## Appendix 2

### Linearization

Stability is established by linearizing about steady-state. We will perform the calculation for the **M** model, then comment on the general form of viable models and their corresponding results. We also use the linearization to demonstrate how simultaneous length control breaks down in the **TM** model. We work out the explicit formulas for the case of diffusive return; the results for ballistic return can be computed with minor changes (see ***Figure 6*** for the resulting formulas).

### M model

Let Δ*L*_1_ and Δ*L*_2_ be the deviations from steady-state such that *L*_1_ = *L*_*ss*_ + Δ*L*_1_ and *L*_2_ = *L*_*ss*_ + Δ*L*_2_. From the Main Text, the flux *J* is given by

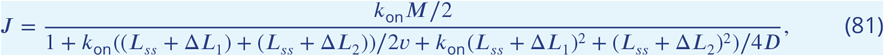

Keeping only those terms in the denominator up to first order in Δ*L*_1_ and Δ*L*_2_,

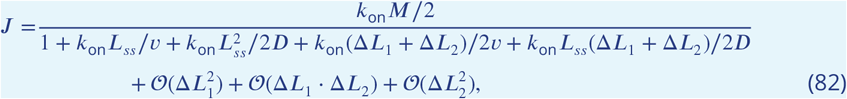

and taking the first-order Taylor series expansion yields

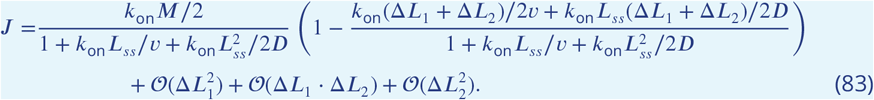

Substituting this expression for the flux into the dynamical equations (12) and (13) we have, to first order,

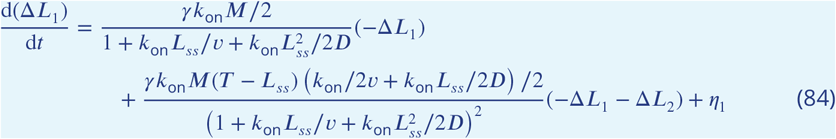

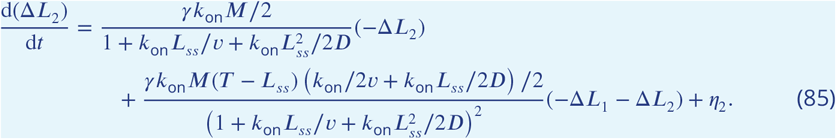

Defining

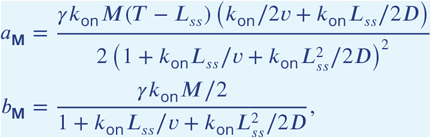

we may write this system in the matrix form

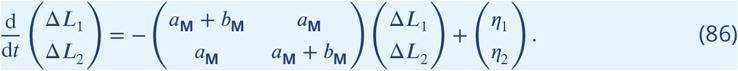

The matrix above is the sum of a rank-1 matrix and a diagonal perturbation. Roughly speaking, *a*_**M**_ corresponds to the shared quantities whereas *b*_**M**_ corresponds to the separate quantities. This system may be diagonalized in terms of the sum ∑ = Δ*L*_1_ + Δ*L*_2_ and difference Γ = Δ*L*_1_ − Δ*L*_2_ to yield

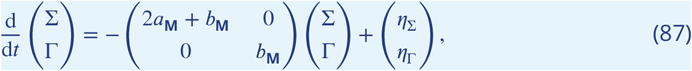

where *η*_∑_ = *η*_Γ_ + *η*_2_ and *η*_Γ_ = *η*_1_ − *η*_2_. Note that the eigenvalues

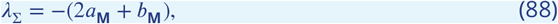

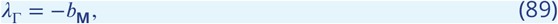

are both negative, so that the steady-state is stable, since *b*_**M**_ is clearly positive and *T* > *L*_*ss*_ implies the positivity of *a*_**M**_. The fact that *λ*_∑_ and *λ*_Γ_ are *distinct* is also important as it provides a possible means to extract two independent parameters from experiment.

### T model

In fact, the matrix form (86) above applies to all 1ve viable models, with suitably modi1ed parameters. For the case of shared tubulin pool and separate motor pools, we have

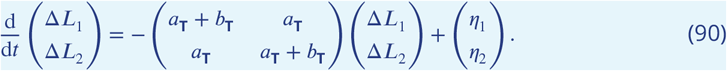

with

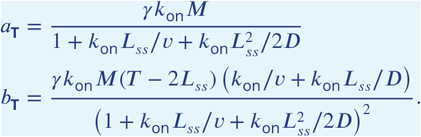

The same logic used in the previous section implies that both eigenvalues are negative and the dynamics is thus stable.

### TM model

Simultaneous length control breaks down if all biomolecules are shared and the disassembly rate is constant. As shown in the Main Text, in this case the steady-state equations provide only a single equation for the two unknowns *L*_1,*ss*_ and *L*_2,*ss*_:

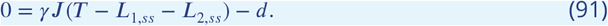

This equation admits infinitely-many solutions *L*_1,*ss*_ and *L*_2,*ss*_. By symmetry, linearizing about any one of these solutions yields a matrix equation of the form

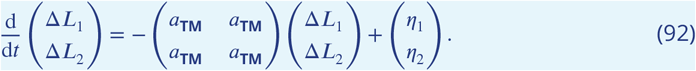

Unlike the analogous matrices for the viable models, the 2 × 2 matrix above has an vanishing eigenvalue, as we now show. Diagonalizing in terms of the sum ∑ = Δ*L*_1_ + Δ*L*_2_ and difference Γ = Δ*L*_1_ − Δ*L*_2_ gives

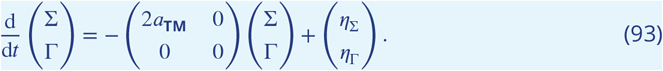

There is a vanishing eigenvalue associated to the difference of lengths, i.e. perturbations from steady-state in the difference of lengths do not decay on a finite timescale. This is consistent with previous results from stochastic simulations that the **TM** model does not yield simultaneous length control ***Mohapatra et al. (2017)***.

### TM* model

For the case of shared tubulin pool and shared motor pools with concentration-dependent disassembly, we have

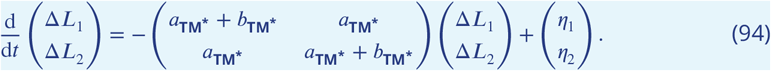

with

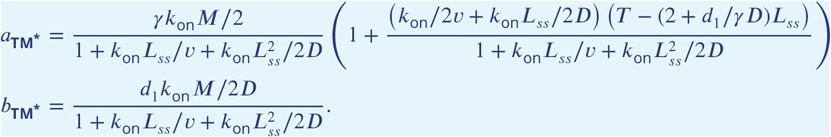

### Arbitrary flagellar number

This matrix form also applies to the case of arbitrary flagellar number *N* discussed in Results. Let Δ*L*_*i*_ := *L*_*i*_ − *L*_*ss*_ for *i* = 1, …, *N* denote the deviation from steady-state for the *i*^th^ flagella. The linearized equations satisfy

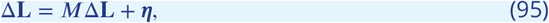

where Δ**L** = (Δ*L*_1_, …, Δ*L*_*N*_), ***η*** = (*η*_1_, …, *η*_*N*_), and the matrix *M* has the form

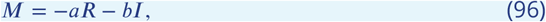

with *R* = **11**^*T*^ the rank-one matrix satisfying *R*_*ij*_ = 1 for all *i, j* and *I* is the *N* × *N* identity matrix, e.g. for N=3

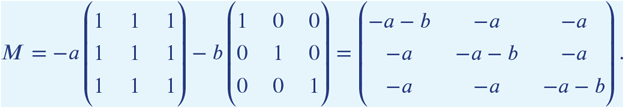

*M* is straightforward to diagonalize. Applying *M* to the vector **v**_1_ := **1** = (1, …, 1)^*T*^ corresponding to the sum of all lengths yields

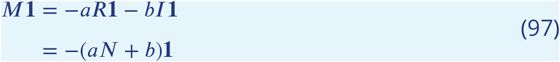

Further, for any vector **x** = (*x*_1_, …, *x*_*N*_) such that 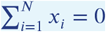, we have *R***x** = 0 and

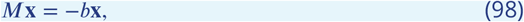

therefore it is an eigenvector with eigenvalue *λ* = −*b*. Note that the pairwise differences

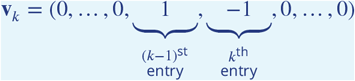

for *k* = 2, …, *N* form a convenient basis for the space of such **x** whose components sum to zero. From (97) and (98) above, in terms of the basis {**v**_1_, **v**_2_, …, **v**_*N*_} consisting of the sum and differences in lengths, the evolution equations diagonalize with eigenvalues

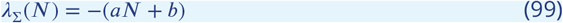

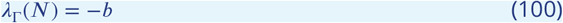

for the sum and differences, respectively, so that *λ*_∑_(*N*) is an eigenvalue of multiplicity 1 and *λ*_Γ_(*N*) is an eigenvalue of multiplicity *N* − 1. Note that in addition to the explicit dependence of *λ*_∑_(*N*) and *λ*_Γ_(*N*) on *N* there is an implicit number-dependence through the steady-state length (and potentially the size of the pool *T*). Since *λ*_∑_(*N*) ≠ *λ*_Γ_(*N*), the autocorrelations described in ***Appendix 3*** will involve two distinct timescales *τ*_∑_(*N*) = *λ*_∑_(*N*)^−1^ and *τ*_Γ_(*N*) = *λ*_Γ_(*N*)^−1^ corresponding to these two eigenvalues.

## Appendix 3

### Fluctuation analysis

We return our attention to the case of *N* = 2 flagella relevant to *Chlamydomonas*. As shown in ***Appendix 2***, linearizing about steady-state results in a diagonal system in terms of the sum ∑ = Δ*L*_1_ + Δ*L*_2_ and difference Γ = Δ*L*_1_ − Δ*L*_2_:

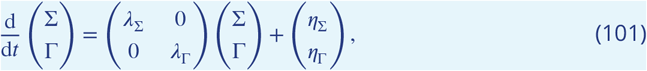

where *η*_∑_ = *η*_1_ + *η*_2_ and *η*_Γ_ = *η*_1_ − *η*_2_ have the covariance structure

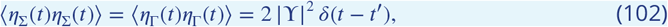

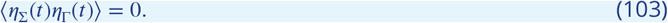

Applying the Fourier transform and solving for 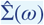 and 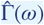 results in a Lorentzian function ***Gardiner (2009)***, as we explicitly show here. We first apply the Fourier transform to (101) to obtain

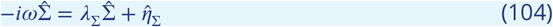

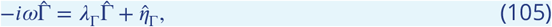

with *λ*_∑_ and *λ*_Γ_ given in ***Appendix 2*** for the viable models. Solving for 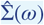 and 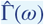 yields

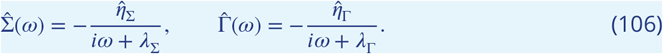

Multiplying by complex conjugates and taking the ensemble average results in

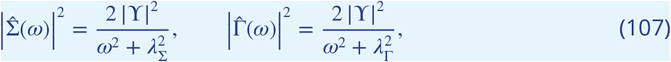

where we have used (102) for the spectra. The Wiener-Khinchin Theorem asserts that for a random variable *X* the autocorrelation function *C*_*X*_(*τ*) := 〈*X*(*t*)*X*(*t* + *τ*)〉 is equal to the inverse Fourier transform of the spectral density gives 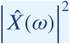. Taking inverse transforms in (107) gives

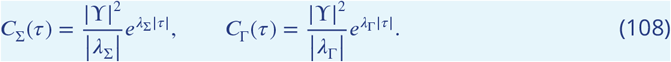

From the linearization in ***Appendix 2***, the predicted timescales based on the parameters in ***Table 1*** are *τ*_∑_ = 17 min and *τ*_Γ_ = 63 min for the **M** model, *τ*_∑_ = 12 min and *τ*_Γ_ = 15 min for the **T** model, and *τ*_∑_ = 17 min and *τ*_Γ_ = 26 min for the **TM\*** model.

### Extracting parameters via fluctuations

As described above, the autocorrelations *C*_∑_(*r*) := 〈∑(*t*)∑(*t* + *τ*)〉 and *C*_Γ_(*τ*) := 〈Γ(*t*)Γ(*t* + *τ*) of the sum and difference in flagellar lengths can be expressed in terms of the timescales *τ*_∑_ := |*λ*_∑_|^−1^ and *τ*_Γ_ := |*λ*_Γ_|^−1^ and the fluctuation magnitudes |ϒ|^2^ *τ*_∑_ and |ϒ|^2^ *τ*_Γ_. These formulas may be used to fit parameters and make testable predictions.

Our models make the prediction *τ*_∑_ < *τ*_Γ_ and |ϒ_∑_|^2^ = |ϒ_Γ_|^2^, which provides a useful test for the overall modeling framework. On a more quantitative level, since *τ*_∑_ ≠ *τ*_Γ_, the autocorrelations provide three distinct predictions, i.e. the formulas for *τ*_∑_, *τ*_r_, and ϒ in terms of model parameters. Experimental measurements can be used to test these predictions (or used to 1t any parameters that cannot be independently estimated). For the concentration-dependent disassembly model **TM**\*, the parameter *d*_1_ is inversely proportional to *τ*_Γ_, making it possible to fit by comparison to the fluctuation spectrum.

To provide a proof of concept, we generate synthetic data using our model **T** to demonstrate that we can use fluctuations to extract the input parameters. We 1t the pooled autocorrelation functions to exponential functions 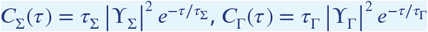 and find that we are able to extract the input values (***Appendix 3 Figure 1(a)-(b)***).

To quantify how much data would be needed to effectively constrain the model parameters, we have performed Monte Carlo simulations with different sampling frequencies and total time durations. ***Appendix 3 Figure 1 (c)-(d)*** shows, for instance, that if the total experiment time is fixed at 1000 minutes (16 hours), sampling intervals of two minutes or less are needed to reliably obtain 10% error in both *τ*_∑_ and *τ*_Γ_. Although the amount of data required is significant, existing protocols are able to maintain healthy cells for over 18 hours using microfluidics ***Ludington et al. (2012)***, so that extracting the model parameters from fluctuations should be experimentally tractable.

**Appendix 3 Figure 1.**
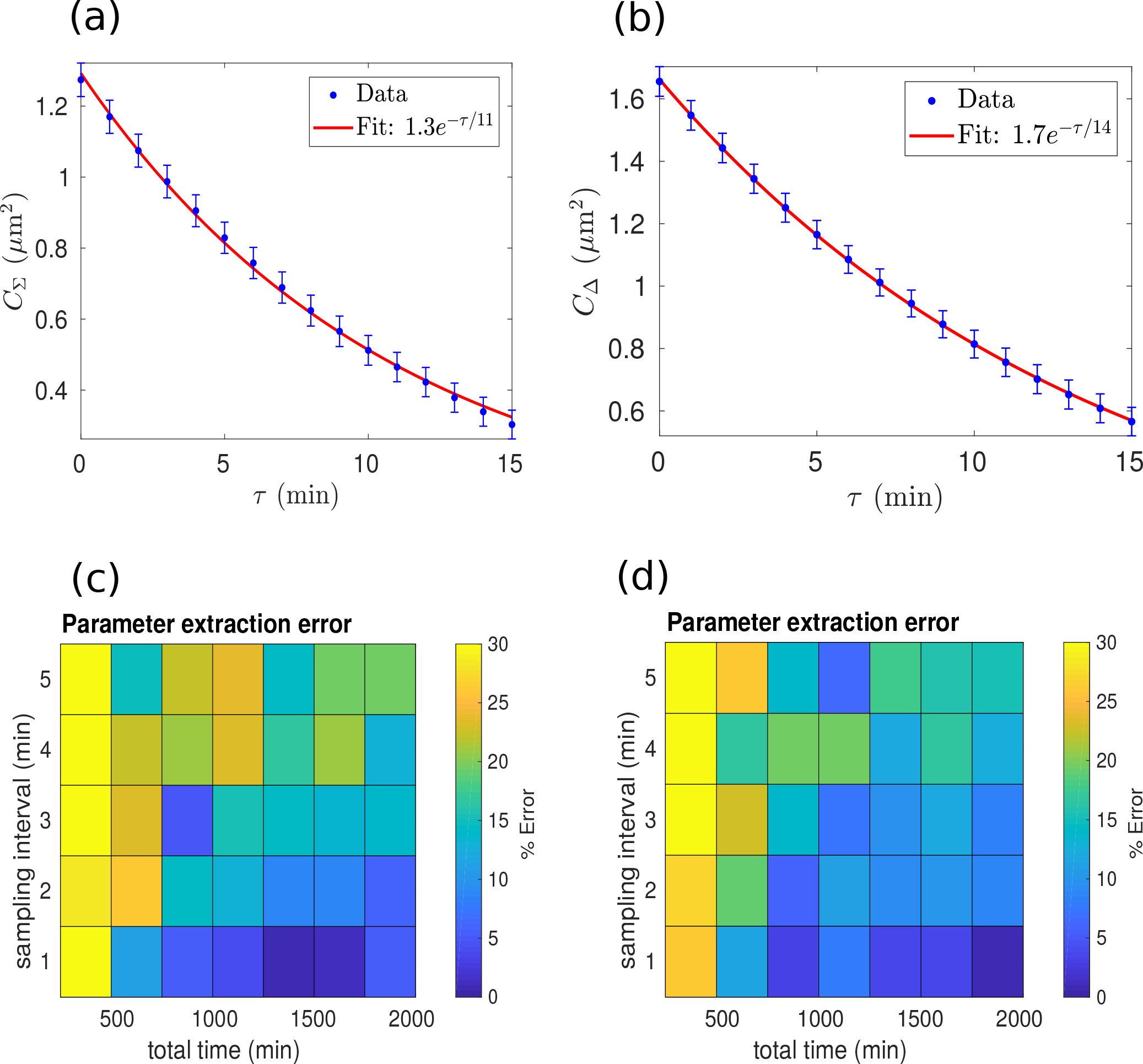
Analysis of synthetic data generated near steady-state using model **T**. The autocorrelations in the sum (a) and difference (b) of flagellar lengths are described by exponential functions with distinct timescales. These timescales can be extracted from data by data fitting. To bound the quantity of data required to accurately extract the timescales, we generated synthetic data for 20 pairs of flagella using different sampling intervals and time durations. We find that using a sampling interval of less than 2 minutes and a time duration of greater than 1000 minutes (approx. 16 hrs) allows the timescales *τ*_∑_ (c) and *τ*_Γ_ (d) to be extracted to within 10% error.

## Footnotes

1 To further justify the condition *c*_*d*_ (0) = 0, consider a microscopic random walk on a one-dimensional lattice with spacing *l*. If the probability of hopping in either direction is equal for each timestep Δ*t*, the flux of motors hopping off the flagellum is 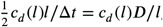, where we have used the relation *D* = *l*^2^ /(2Δ*t*) between the continuous diffusion coeZcient and parameters of the discrete random walk. For the fluxes onto and off of the flagellum to balance, 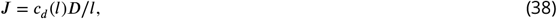 so that in the limit *l* → 0 we must have *c*_*d*_ (0) = 0 in order to obtain finite *J*.

